# Propofol Reverses Surgery-Induced Neuroinflammation and Cognitive Impairment in Aged Mice via α5-GABA_A_ Receptors

**DOI:** 10.1101/2022.10.26.513964

**Authors:** Rajasekar Nagarajan, Jinrui Lyu, Maltesh Kambali, Muxiao Wang, Robert A. Pearce, Uwe Rudolph

**Affiliations:** Department of Comparative Biosciences, College of Veterinary Medicine; Neuroscience Program; Carl R. Woese Institute for Genomic Biology, University of Illinois Urbana-Champaign, Urbana, Illinois, USA; Department of Anesthesiology, University of Wisconsin-Madison, Madison, WI, USA

**Keywords:** Propofol, Surgery, α5-GABA_A_ Receptors, Perioperative Neurocognitive Disorder, Anesthesia, Microglial Activation

## Abstract

Surgery may lead to long-lasting cognitive deficits that are referred to as perioperative neurocognitive disorder (NCD), particularly in elderly patients. Currently, no interventions are routinely employed in clinical practice to prevent perioperative NCD. Here we show that perioperative chronic intermittent administration of propofol to aged mice undergoing laparotomy under isoflurane anesthesia effectively blocks the surgery-induced increase in nitrosative stress, increased expression of proapoptotic proteins, microglial activation, and cognitive deficits. By contrast, in the absence of surgery and anesthesia, propofol had little effect on biochemical parameters and led to cognitive improvement only in a subset of behavioral paradigms. The actions of propofol were largely absent in mice lacking the GABA_A_ receptor α5-subunit, indicating that they are mediated by α5-containing GABA_A_ receptors. These results demonstrate that propofol – via α5-containing GABA_A_ receptors that are redistributed to the cell surface membranes in a sustained manner – can attenuate surgery-induced neuroinflammation and postsurgical cognitive deficits.

## 1. Introduction

Cognitive decline after surgery which can persist for months has been referred to as postoperative cognitive dysfunction (POCD)(1, 2) or more recently as perioperative neurocognitive disorder (NCD) (3). It is observed mainly in elderly patients. In patients >60 years of age, approximately 26% of patients display signs of cognitive dysfunction one week after surgery, and 10% of patients still show signs three months after surgery, compared to 3% in controls at both time points(1) There is no evidence in the clinical literature that general anesthetics themselves cause NCD, as comparisons of regional versus general anesthesia have not found differences in NCD rates; thus, it is likely that other factors related to surgery are important(4). Indeed, there is evidence that postoperative deficits in mice are due to surgery itself, which induces an inflammatory response(5), but not the general anesthetic, e.g., isoflurane (6). In addition, Xu et al. (2014) reported that 18- month-old but not 9-month-old mice display cognitive impairments after a laparotomy under local anesthesia, accompanied by an increase in β-amyloid that is attenuated by a γ-secretase inhibitor, suggesting that aged mice are more sensitive to the cognition-impairing effects of surgery(7)

Propofol (2, 6-diisopropyl phenol) is a potent intravenous hypnotic agent widely used for induction and maintenance of anesthesia during surgery. Apart from its general anesthetic effects, it has been reported to possess anti-oxidative effects (8), anti-inflammatory potential (9) and neuroprotective properties (10, 11). Furthermore, Shao et al. (2014) have shown that chronic intermittent treatment with propofol in aged rodents improves cognitive functions and attenuates caspase-3 activation (12) In addition, propofol has been reported to protect against the irreversible neurodegenerative reaction induced by the NMDA antagonist, MK-801(13). In contrast, multiple exposures of propofol induced persistent neuronal apoptosis, neuronal deficit, synaptic loss, and long-term cognitive impairment in neonatal rodents (14). The pharmacological mechanisms underlying the various actions of propofol (from neuroprotection to neurotoxicity) and for the finding that chronic intermittent administration of propofol increases cognitive performance in aged mice and in a transgenic mouse model of Alzheimer’s Disease (AD Tg)(12) are unknown.

GABA_A_ receptors are recognized as major targets for the general anesthetic action of intravenous anesthetics such as propofol and etomidate(15). The α5-GABA_A_ receptors play an important role in controlling hippocampal-dependent memory under physiological conditions(16, 17) and have been shown to mediate the acute amnestic actions of etomidate both in the acquisition phase and the recall phase(18). In addition to its canonical acute amnestic actions, etomidate has also been shown to have non-canonical sustained amnestic actions for up to 7 days, when the drug is likely no longer present in the animal, which are due to a redistribution of α5-GABA_A_ receptors to the cell surface membranes(19). An increased cell surface expression of α5-GABA_A_ receptors has also been observed 24 h after an anesthetizing dose of isoflurane(19). Negative allosteric modulation of α5-GABA_A_ receptors by pretreatment with L-655,708 has been shown to reverse the amnestic effects of etomidate and isoflurane(20, 21), which led to the idea that α5-negative allosteric modulators might be suitable to prevent or to treat postoperative cognitive deficits.

The experiments by Zurek et al. (2014) reporting an increase in surface expression of α5-GABA_A_ receptors, a sustained increase in tonic inhibition in CA1 pyramidal neurons and sustained cognitive impairments after etomidate (19) are consistent with the prevailing view that α5-GABA_A_ receptors would primarily impair cognitive functions. They were done in 3- to 5-month-old mice. In contrast, it has been shown that in aged rats (but not in young adult rats) positive allosteric modulation of α5-GABA_A_ receptors leads to a cognitive improvement (22), suggesting that the role of α5-GABA_A_ receptors in modulation of cognition may be age-dependent. As NCD is a problem primarily in aged individuals, we decided to perform our studies on postoperative cognitive deficits in aged mice (21-24 months of age). Our hypothesis was that in aged mice undergoing laparotomy perioperative chronic intermittent propofol would lead to a redistribution of α5-GABA_A_ receptors to the cell surface membranes, and that this increase in surface α5-GABA_A_ receptors would reverse postoperative cognitive deficits. Moreover, while a role of α5-GABA_A_ receptors in surgery-induced nitrosative stress, neuroinflammation and apoptosis was unknown to date, we hypothesized that an increased availability of α5-GABA_A_ receptors would counteract these surgery-induced changes, which might contribute to a prevention or reversal of surgery-induced cognitive dysfunction.

## 2. Results

### 2.1. Propofol induces a sustained redistribution of α5-GABA_A_ receptors to cell-surface membranes in the hippocampus

To determine whether propofol affects the amount of α5-GABA_A_ receptors in the cell surface membranes, we performed biotinylation experiments followed by Western blots. We determined both surface α5 protein expression and total α5 protein expression after propofol treatment (100 mg/kg i.p.) at different time points: 24h, 72h, five days, and seven days. We found increased cell-surface expression of α5-GABA_A_ receptor in hippocampus after 24h (*p <0.05), 72h (*p<0.05), and five days (*p< 0.05), but not at seven days (p>0.05) after treatment with propofol; expression of the **α**5 subunit in total membranes was unchanged (p>0.05) (Fig 1). Further, we studied the effect of chronic intermittent propofol (CIP), specifically 75 mg/kg, i.p., every 5^th^ day for 21 days on α5-GABA_A_ receptor expression in surface and total membrane. We found that CIP treatment for 21 days significantly up-regulated the levels of cell-surface expression of α5 subunits (*p<0.05), with no change of the amount of the α5 subunit in total membranes (p>0.05) (Fig. 2A,B). To confirm the sustained redistribution of α5-GABA_A_ receptors to cell-surface membranes in aged mice (21-24 months old), propofol was injected at a dosage of 75 mg/kg intraperitoneally. Consistent with the findings in young mice, increased cell-surface expression of α5-GABA_A_ receptors was observed in the hippocampus after five days (*p < 0.05, 2C), whereas the expression of the **α**5 subunit in total membranes was unchanged (p>0.05, 2D). Overall, the results indicate that propofol induces a sustained redistribution of α5- GABA_A_ receptors to cell-surface membranes in the hippocampus. This effect was observed in both young adult and aged mice, and with repeated propofol administration indicating that there is no development of tolerance in terms of receptor redistribution to the cell surface membranes.

**Figure 1.**
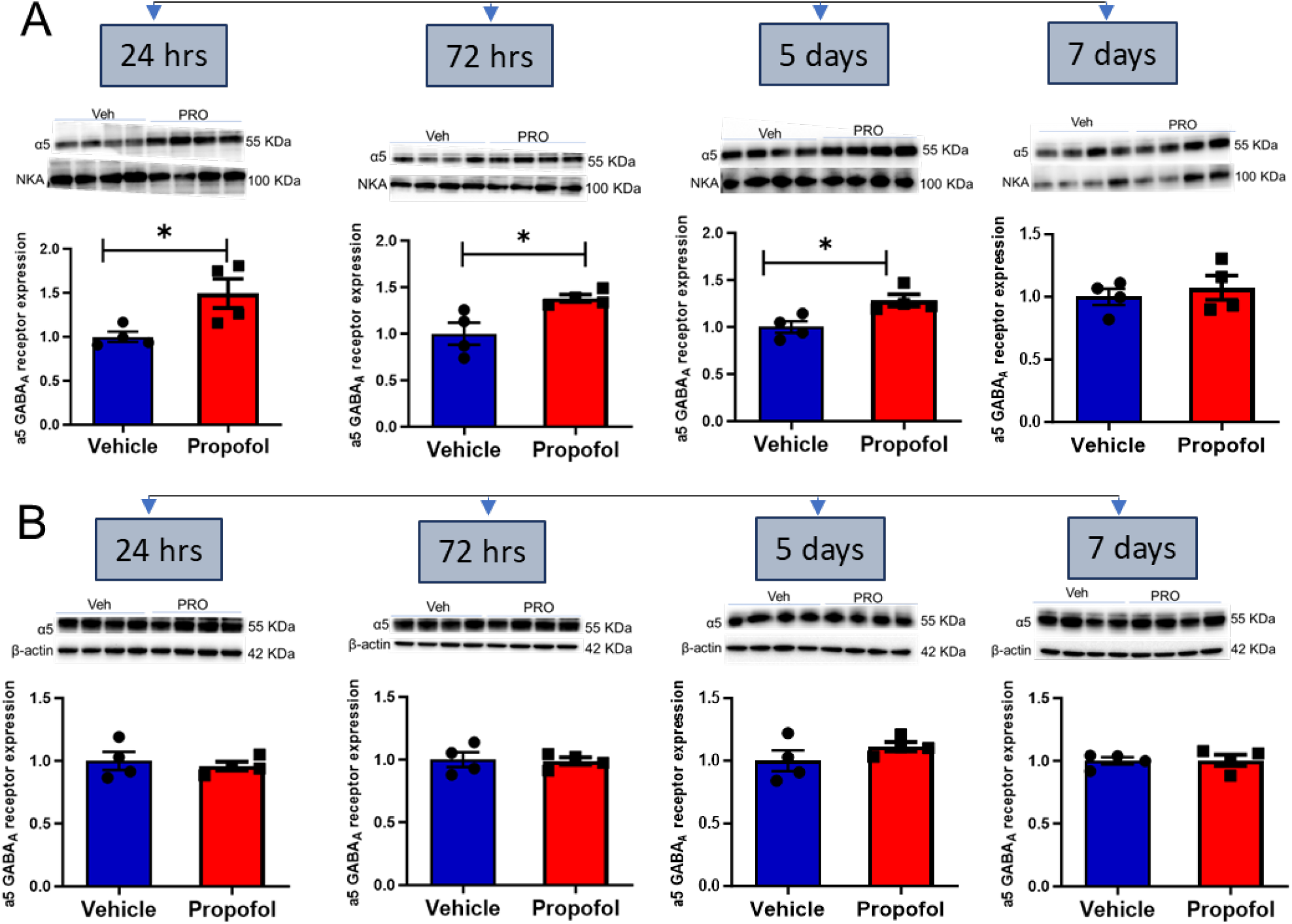
Time course of cell surface expression of α5-GABA_A_ receptors in hippocampus after a single dose of propofol. Biotinylation was performed followed by Western blots in young adult mice. The α5-GABA_A_ receptor expression in cell surface membranes was normalized with Na+/K+ ATPase (NKA) (**A**), and the α5-GABA_A_ receptors expression on total membranes was normalized with β-actin (**B**). Bar graph images depict the mean ± S.E.M. of the relative density of protein (n=4). Unpaired two-tailed t-tests were conducted to analyze the data. Significance (*) indicates a significant difference (*p < 0.05) compared to the vehicle control. The molecular weight (MW) is represented in kilodaltons (kDa). For Figure 1A, the statistical results are as follows: 24 hours: t(6)=2.8, p=0.0310; 72 hours: t(6)=3.1, p=0.0224;5 days: t(6)=3.2, p=0.0185;7 days: t(6)=0.63, p=0.5513. For Figure 1B, the statistical results are as follows:24 hours: t(6)=0.51, p=0.6256; 72 hours: t(6)=0.18, p=0.8645; 5 days: t(6)=1.3, p=0.2575;7 days: t(6)=0.10, p=0.9201.

**Figure 2.**
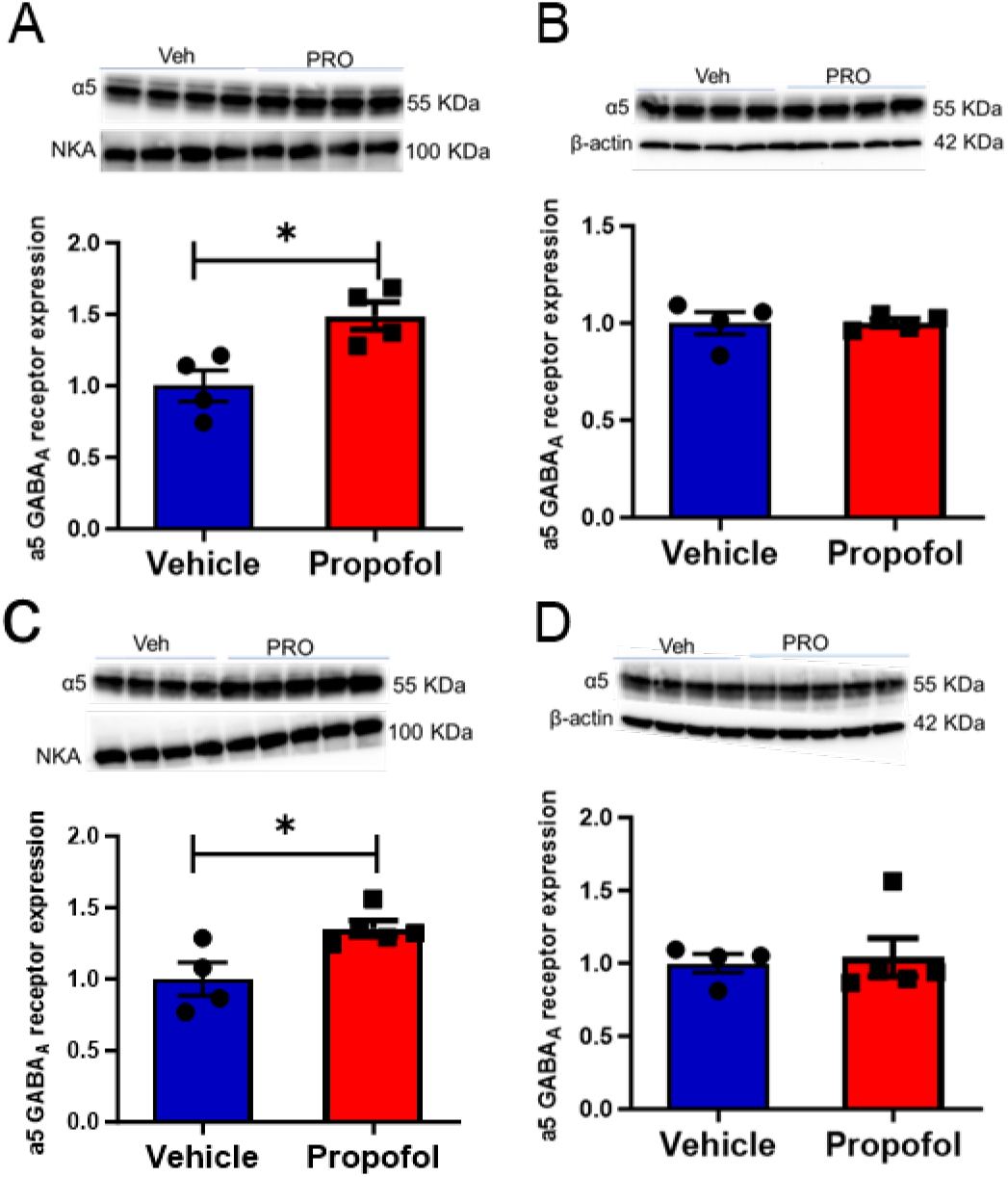
Cell surface expression of α5-GABA_A_ receptors after chronic intermittent propofol administration in young adult mice and after single dose propofol administration in aged mice in the hippocampus. Young adult mice were subjected to either chronic intermittent propofol (CIP) or vehicle (Intralipid^R^) administration every 5^th^ day for a total of 21 days. The expression of α5-GABA_A_ receptors on the cell surface membrane was normalized using Na+/K+ ATPase (NKA) (**A**), while the expression of α5-GABA_A_ receptors on total membranes was normalized using β- actin (**B**). In aged mice, cell surface expression of α5-GABA_A_ receptors in the hippocampus was studied following a single dose of propofol or vehicle (Intralipid^R^) administration. Biotinylation was performed 5 days after the propofol administration, followed by Western blot analysis. The expression of α5-GABA_A_ receptors on cell surface membranes was normalized using Na+/K+ ATPase (NKA) (**C**), and the expression of α5-GABA_A_ receptors on total membranes was normalized using β-actin (**D**). The bar graph images illustrate the mean ± S.E.M. of the relative protein density (n=4-5). The molecular weight (MW) is displayed in kilodaltons (kDa). Unpaired two-tailed t-tests were performed to analyze the data, with significance (*) indicating a significant difference (*p < 0.05) compared to the vehicle control. The statistical results are as follows: A: t(6)=3.4, p=0.0152; B: t(6)=0.035, p=0.9734; C: t(7)=3, p=0.0204; D: t(7)=0.26, p=0.8040.

### 2.2. Examination of the effects of CIP treatment on surgery-induced behavioral and biochemical changes

To evaluate the impact of CIP treatment during the perioperative phase and to explore the potential involvement of α5-GABA_A_ receptors in the hypothesized protective response mediated by CIP, we implemented the experimental protocol outlined in Figure 3. WT or α5 knockout (α5 KO) mice received 75 mg/kg propofol i.p. every 5 days throughout the experiment starting on day 1. Laparotomy was performed on day 17, followed by open field test on day 19, Y maze test on day 20, novel object recognition (NOR) test on day 21-22, fear conditioning on days 23-25 (day 23: training; day 24: contextual test; day 25: tone test), Morris water maze on days 26-39, and fear conditioning testing on days 40 (contextual test) and 41 (tone test). Mice were euthanized for molecular analysis on day 42 (Fig. 3).

**Figure 3.**
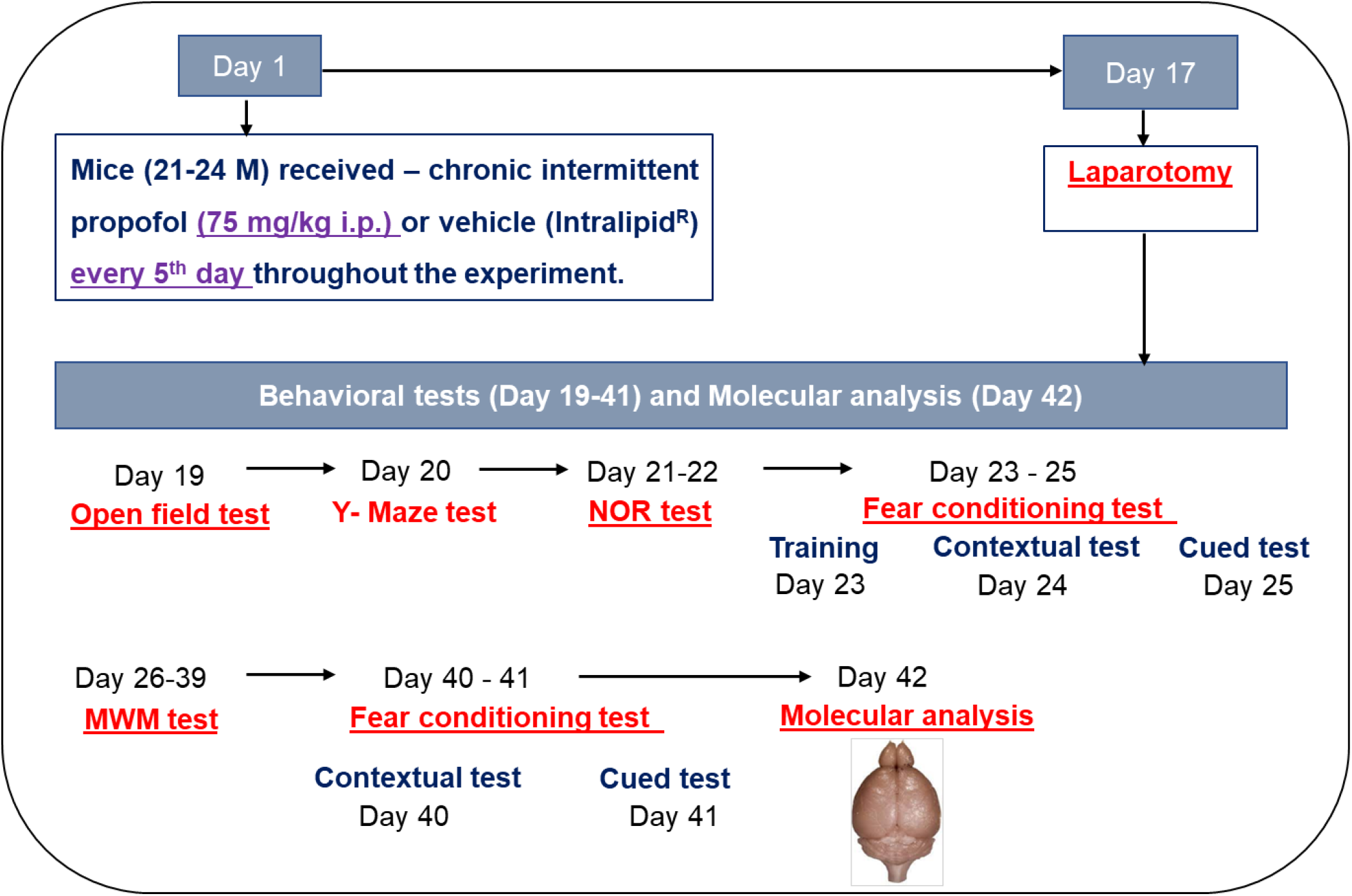
A schematic overview of the experimental protocol designed to investigate the effects of perioperative chronic intermittent propofol (CIP) treatment. The study involves two groups of mice: wild type C57BL/6J mice and α5 knockout mice. NOR - Novel object recognition; MWM - Morris water maze.

#### 2.2.1. Open field test

We assessed the effects of perioperative CIP treatment and/or laparotomy surgery on open-field behavior in aged WT and α5 KO mice (21-24 months old), specifically on locomotor activity and anxiety-like behavior. Our findings indicate that neither the CIP treatment nor the surgery had a significant impact on the total distance traveled, compared to the vehicle control condition, in both WT (Fig. 4A) and α5 KO mice (Fig. 4C). Moreover, the CIP treatment did not result in any significant changes in the time spent in the center of the open field for the WT mice (Fig. 4B). However, the surgery significantly reduced the time spent in the center of the open field for the α5 KO mice (p<0.05), indicative of anxiety-like behavior (Fig. 4D). Notably, the CIP treatment led to a significant increase, of the inner zone cumulative duration in the open field test compared to the Surgery + vehicle group in α5 KO mice (p<0.05), i.e., to a reversal of the surgery-induced anxiety-like behavior (Fig. 4D). This indicates that the α5 KO mice are more susceptible to express anxiety-related behavior after surgery, which is in line with previous findings showing that α5- GABA_A_ receptors may modulate anxiety-like behaviors (23, 24). Taken together, our data suggest that both the CIP treatment and the surgery did not disrupt open field behavior in WT mice or total distance moved in the α5 KO mice, indicating that CIP treatment does not have an effect on locomotion.

**Figure 4.**
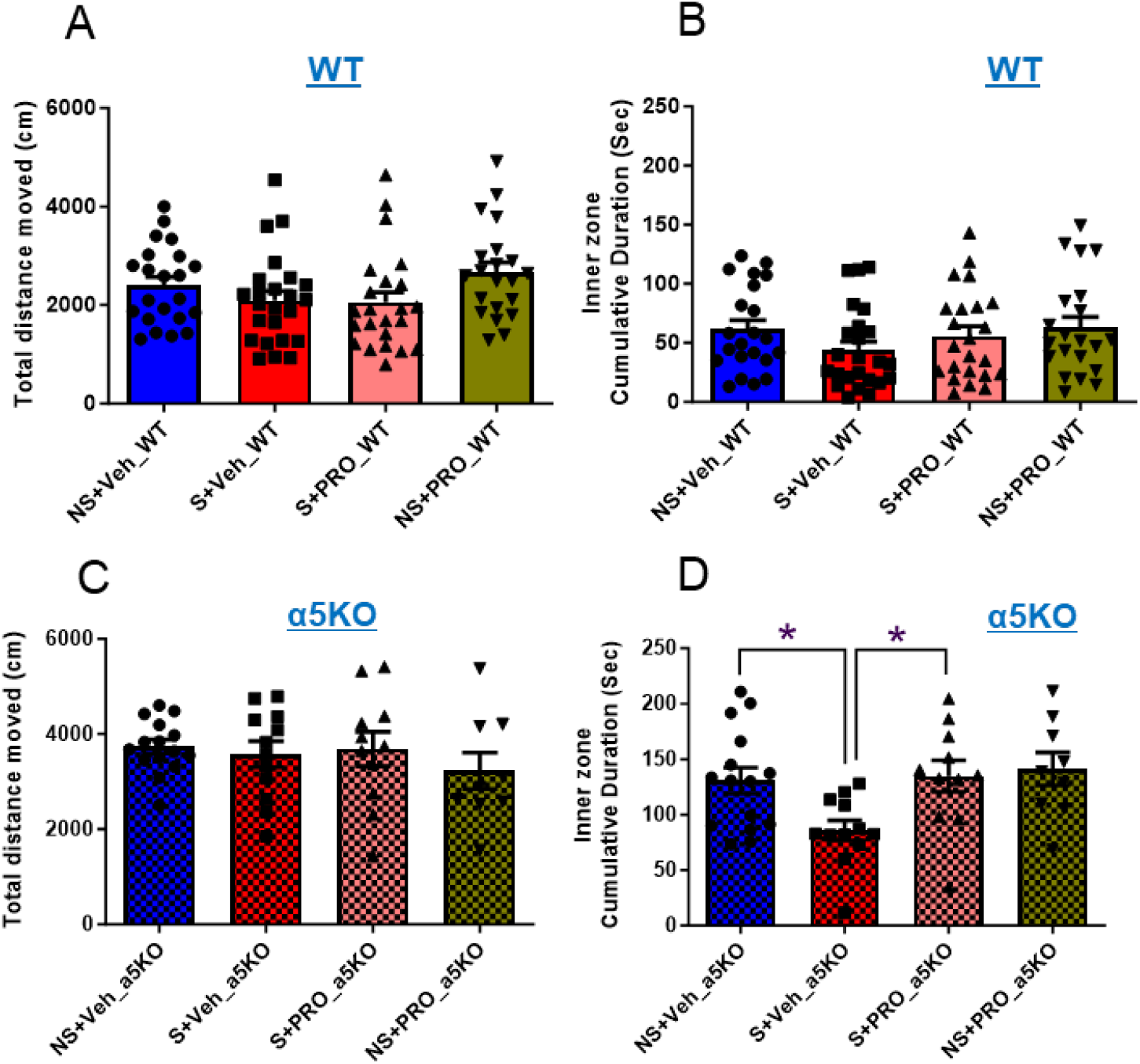
The effects of surgery and chronic intermittent propofol (CIP) treatment on the open field test in aged wild type and α5 knockout mice. The following parameters were measured: **A**) Total distance moved (in centimeters), **B**) Cumulative duration in the inner zone (in seconds) for wild type mice. **C**) Total distance moved (in centimeters), **D**) Cumulative duration in the inner zone (in seconds) for α5 knockout mice. The results are presented as mean ± SEM. The abbreviations used are as follows: NS+Veh: no surgery control + vehicle; S+Veh: surgery + vehicle; S+PRO: surgery + propofol; NS+PRO: no surgery control + propofol; WT: wild type mice; α5KO: α5 knockout mice. Significance (*) indicates a significant difference (*p < 0.05). Statistical analyses were performed using GraphPad Prism, employing 1-way ANOVA followed by Tukey’s test for multiple comparisons. The results are as follows: A) F (3, 86) = 2.099, P=0.1063; B) F (3, 86) = 1.174, P=0.3244; C) F (3, 43) = 0.5931, P=0.6229; D) F (3, 43) = 4.049, P=0.0128 (NS+Veh vs S+Veh, q=3.86; S+Veh vs S+PRO, q=3.901).

#### 2.2.2. Y maze and novel object recognition (NOR) tests

In the Y maze test, surgery resulted in a significant reduction in the percentage of alternations in both aged WT (p<0.05) (Fig. 5A) and aged α5 KO mice (p<0.05) (Fig. 5C). Conversely, CIP treatment significantly increased the percentage of alternations in the Y maze test compared to the no-surgery control and surgery control groups (p<0.05) (Fig. 5A), but only in aged WT mice and not in aged α5 KO mice (p>0.05) (Fig. 5C). In the novel object recognition test, surgery led to a reduction in the recognition index compared to the non-surgery control group in both aged WT (p<0.05) (Fig. 5B) and α5 KO mice (p<0.05) (Fig. 5D). Remarkably, CIP treatment reversed the surgery-induced reduction of the recognition index in the NOR test (p<0.001) (Fig. 5B), but again, this effect was observed only in aged WT mice (p<0.05) (Fig. 5B) and not in aged α5 KO mice (p>0.05) (Fig. 5D). No significant differences were observed between males and females in the Y maze and NOR tests (data not shown). These findings indicate that CIP treatment can reverse the impairments of working memory induced by surgery in the Y maze test and NOR test. Furthermore, our results suggest that CIP treatment alone, in the absence of surgery, is sufficient to improve performance in these tests in wild type mice. The absence of these effects in α5 KO mice indicates that they are mediated by the α5 GABA_A_ receptor.

**Figure 5:**
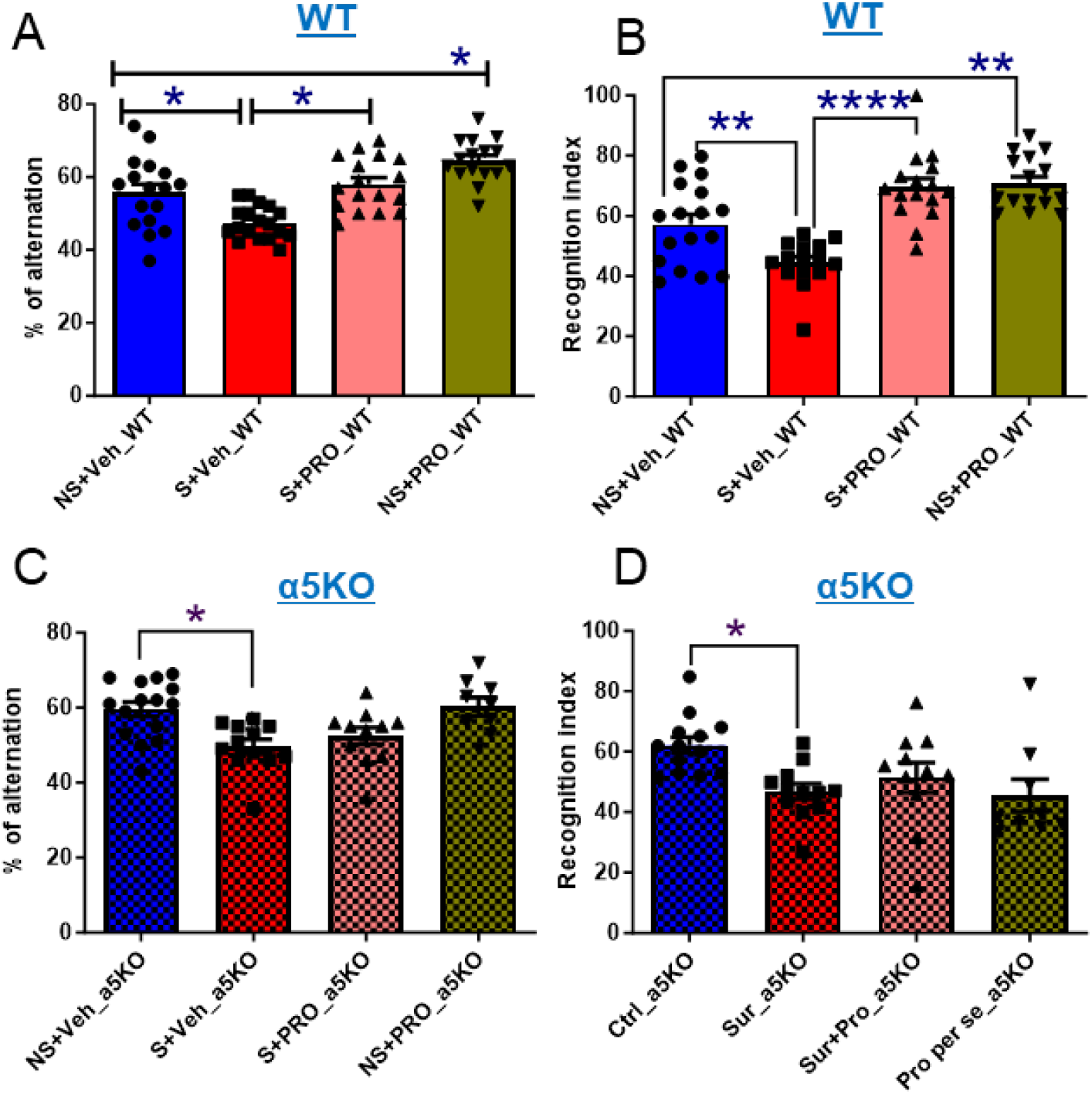
Effects of surgery and CIP on the Y maze test and the novel object recognition (NOR) test in aged wild type (A, C) and α5 knockout mice (B, D). **A**) Spontaneous arm alternation percentage in the Y-maze test for wild type mice, F (3, 64) = 17.07, P<0.0001 (comparisons: NS+Veh vs S+Veh, q=4.951; S+Veh vs S+PRO, q=6.308, NS+Veh vs NS+PRO, q=5.014). **B**) Recognition index in the novel object recognition (NOR) test for wild type mice, F (3, 60) =20.33, P<0.0001 (comparisons: NS+Veh vs S+Veh, q=4.737; S+Veh vs S+PRO, q=9.172, NS+Veh vs NS+PRO, q=5.143). **C**) Spontaneous arm alternation percentage in the Y-maze test for α5 knockout mice, F (3, 43) = 6.091, P=0.0015 (comparison: NS+Veh vs S+Veh, q=4.971). **D**) Recognition index in the novel object recognition (NOR) test for α5 knockout mice, F (3, 41) = 4.124, P=0.0121 (comparison: NS+Veh vs S+Veh, q=4.217). Results are expressed as mean ± SEM and statistically evaluated by one-way analysis of variance followed by Tukey’s Multiple Comparison Test.*p<0.05, **p<0.01, ***p<0.001. The abbreviations used in the figure are as follows: NS+Veh: no surgery control + vehicle. S+Veh: surgery + vehicle. S+PRO: surgery + propofol. NS+PRO: no surgery control + propofol. WT: wild type mice. α5KO: α5 knockout mice.

#### 2.2.3. Morris water maze (MWM) in WT mice

In the MWM, there were significant differences in escape latency (p<0.05, Fig. 6A) and path length (p<0.05, Fig. 6B) in the WT mice that have undergone surgery compared to control mice without surgery in both learning phase (Day 1-8) (Fig. 6A,B) and reversal learning phase (Day 10- 13) (Fig. 6A,B). Surgery led to an increase in the escape latency (Fig. 6A) and path length (Fig. 6B) in both learning and reversal learning phases. CIP treatment decreased the escape latency (p<0.05, Fig. 6A) and path length (p<0.05, Fig. 6B) in the surgery group (Surgery + Propofol) as compared to the vehicle-treated group (Surgery + Vehicle) in both learning phase (Day 1-8, Fig. 6A and 6B) and reversal learning phase (Day 10-13, Fig. 6A and 6B), thus reversing the effects induced by surgery. The treatment with CIP did not significantly alter the escape latency (p>0.05, Fig.6A) and path length (p>0.05, Fig.6B) in non-surgery control mice. There were no significant differences in swimming speed between the surgery, non-surgery controls, and CIP-treated groups (data not shown).

**Figure 6:**
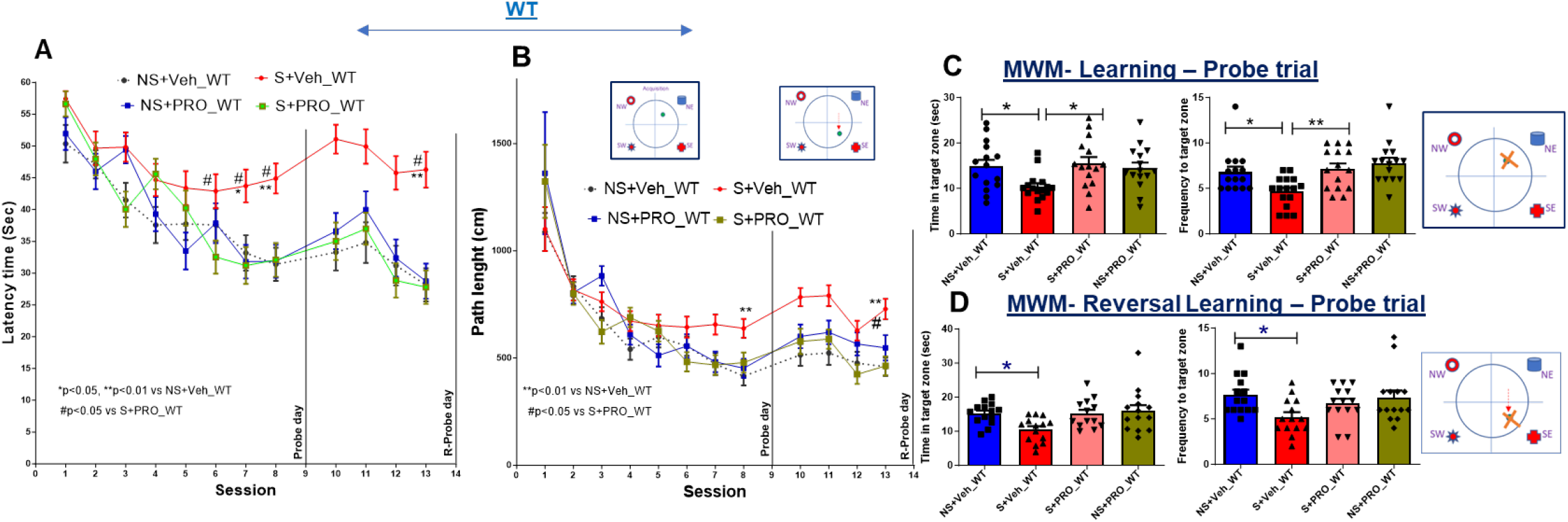
Effects of surgery and chronic intermittent propofol in the Morris water maze task in aged wild type mice. The Morris water maze consists of a learning phase with sessions 1-8 (day 1-8) and learning phase probe trials in session 9 (Day 9). It also includes a reversal learning phase with sessions 10-13 (day 10-13) and reversal learning probe trials on day 14. **A**) Latency time (in seconds) to reach the hidden platform. **B**) Path length (in centimeters) to the hidden platform. Statistical significance: *p<0.05, **p<0.01 (S+Veh vs NS+Veh); #p<0.05 (S+Veh vs S+PRO). **C**) Cumulative duration (Time in target zone) F (3, 57) = 3.826, P=0.0145 (comparisons: NS+Veh vs S+Veh, q= 3.775; S+Veh vs S+PRO, q= 4.264) and frequency of entering the target platform area, F (3, 57) = 6.387, P=0.0008 (comparisons: NS+Veh vs S+Veh, q= 3.960; S+Veh vs S+PRO, q= 4.657) in the learning phase probe trials. **D**) Cumulative duration (Time in target zone), F (3, 51) = 4.269, P=0.0092 (comparisons: NS+Veh vs S+Veh, q= 3.895) and frequency of entering the target platform area, F (3, 52) = 3.148, P=0.0327 (comparisons: NS+Veh vs S+Veh, q= 3.976) in the reversal learning phase probe trials. Statistical significance: *p<0.05, **p<0.01. Abbreviations: NS+Veh: no surgery control + Vehicle. S+Veh: surgery + vehicle. S+PRO: surgery + propofol. NS+PRO: no surgery control + propofol. WT: wild type mice. Statistical analyses were performed using GraphPad Prism. Panels A and B were analyzed with Two-way ANOVA followed by Sidak’s multiple comparisons test, while Panels C and D were analyzed using 1-way ANOVA followed by Tukey’s test for multiple comparisons.

Probe trials were conducted to assess spatial reference memory after both learning phase (learning phase probe trials) (Day 9), and reversal learning phase (reversal learning probe trials) (Day 14). Cumulative duration (Time in target zone) and frequency to enter the target platform area in learning phase probe trials and reversal learning probe trials were recorded. In the learning phase probe trials, surgery reduced both time in the target zone (p<0.05) (Fig. 6C) and frequency to enter the target zone (p<0.05) (Fig. 6C) compared with no-surgery controls. Likewise, in the reversal learning phase probe trials surgery also reduced time in the target zone (p<0.05) (Fig. 6D) and frequency to enter the target zone (p<0.05) (Fig. 6D) compared with the no-surgery control group. The results indicate that surgery impaired reference memory. CIP treatment of mice that underwent surgery significantly increased time in the target zone (p<0.05, Fig 6C) and frequency of entering the target zone (p<0.05, Fig. 6C) in the learning phase probe trial but not in the reversal learning probe trials (p>0.05, Fig 6D & p>0.05, Fig 6D). Our findings indicate that surgery impairs learning and reversal learning, and that this impairment can be reversed by CIP in the learning phase.

#### 2.2.4. Morris water maze (MWM) in a5 KO mice

In α5 KO mice, significant differences were observed in escape latency (p<0.05, Fig. 7A) and path length (p<0.05, Fig.7B) between mice that underwent surgery and control mice without surgery in both the learning phase (Day 1-6) and the reversal learning phase (Day 8-11). Surgery led to increased escape latency (Fig. 7A) and path length (Fig. 7B) during both learning and reversal learning phases. However, treatment with CIP did not decrease the escape latency (p>0.05, Fig. 7A) and path length (p>0.05, Fig. 7B) in the surgery group (Surgery + Propofol) compared to the vehicle-treated group (Surgery + Vehicle) in both learning and reversal learning phases. Additionally, CIP treatment did not significantly alter the escape latency (p>0.05, Fig. 7A) and path length (p>0.05, Fig. 7B) in the no-surgery control mice. There were no significant differences in swimming speed between the surgery, no-surgery control, and propofol-treated groups (data not shown).

**Figure 7.**
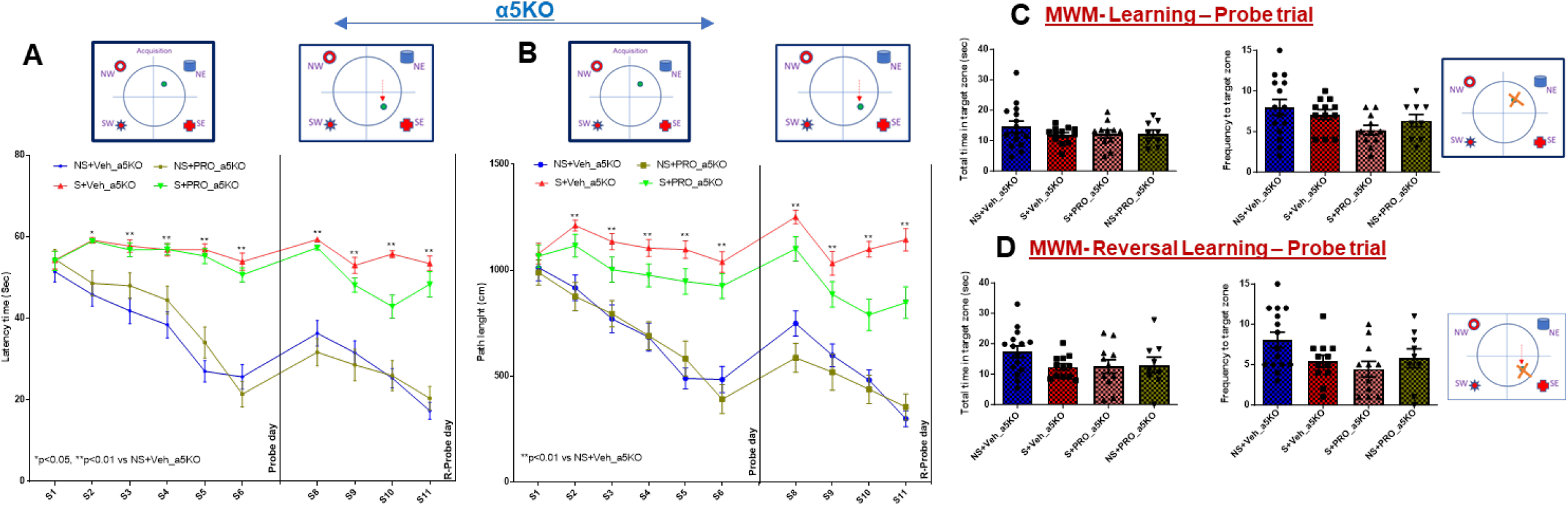
Effects of surgery and chronic intermittent propofol in the Morris water maze task in aged α5 knockout mice. The Morris water maze consists of a learning phase with sessions 1-6 (day 1-6) and learning phase probe trials in session 7 (Day 7). It also includes a reversal learning phase with sessions 8-11 (day 8-11) and reversal learning probe trials on day 12. **A**) Latency time (in seconds) to reach the hidden platform. **B**) Path length (in centimeters) to the hidden platform. Statistical significance: *p<0.05, **p<0.01 (S+Veh vs NS+Veh). **C**) Cumulative duration (time in target zone) and frequency of entering the target platform area in the learning phase probe trials. **D**) Cumulative duration (time in target zone) and frequency of entering the target platform area in the reversal learning phase probe trials. Abbreviations: NS+Veh: no surgery control + Vehicle. S+Veh: surgery + vehicle. S+PRO: surgery + propofol. NS+PRO: no surgery control + propofol. α5KO: α5 knockout mice. Statistical analyses were performed using GraphPad Prism. Panels A and B were analyzed with Two-way ANOVA followed by Sidak’s multiple comparisons test, while Panels C and D were analyzed using 1-way ANOVA followed by Tukey’s test for multiple comparisons.

Probe trials were conducted to assess spatial reference memory after the learning phase (learning phase probe trials) (Day 7) and the reversal learning phase (reversal learning probe trials) (Day 12). Cumulative duration (time in target zone) and frequency of entering the target platform area were recorded. In the learning phase probe trials, time to the target zone (Fig. 7C) and frequency of entering the target zone (Fig. 7C) were not significantly changed in the surgery group compared to the no-surgery control group, (F (3, 43) = 0.8192, P=0.4904, q=1.921 for time in target zone; F (3, 43) = 2.340, P=0.0867, q=1.215 for frequency of entering the target zone). Likewise, during the reversal learning phase probe trials, surgery did not result in a significant change in the time spent within the target zone (as depicted in Fig. 7D). However, it notably led to a reduction in the frequency of entries into the target zone (also shown in Fig. 7D) [(F (3, 43) = 1.714, P=0.1782, q=2.688 for time in target zone; F (3, 43) = 2.864, P=0.0477, q=2.920 for frequency of entering the target zone)].These results suggest that surgery did not impair reference memory in the α5 KO mice. Moreover, CIP treatment in α5 KO mice that have undergone surgery did not significantly increase time in the target quadrant or frequency of entering the target platform area in the learning phase probe trial (p>0.05 and p>0.05, Fig. 7C) or reversal learning probe trials (p>0.05 and p>0.05, Fig. 7D). Also, CIP had no effect in the absence of surgery in α5 KO mice in both learning probe trials (p>0.05 and p>0.05, Fig. 7C) and reversal learning probe trails (p>0.05 and p>0.05, Fig. 7D).

Thus, our findings demonstrate that surgery impairs learning and reversal learning, and that this impairment can be reversed by CIP in WT mice but not in α5 KO mice, indicating that CIP reverses surgery-induced impairments in spatial memory via α5-GABA_A_ receptors.

#### 2.2.5. Trace and contextual fear conditioning

Fear conditioning is among the most commonly used behavioral tests to detect cognitive impairment induced by anesthesia or anesthesia plus surgery (7, 25). In aged wild type mice, we observed a reduction in freezing time in both the tone test (p<0.05, Fig 8A,E) and the context test (p<0.05, Fig 8B,F) at both 1-2 days (Fig. 8A,B) and 17-18 days (Fig. 8E,F) after training in the surgery group compared to non-surgery controls. However, treatment with CIP significantly increased freezing, i.e., reversed the surgery-induced reduction of freezing, in the tone test (p<0.05, Fig 8A) and the context test (p<0.05, Fig 8B) at 1-2 days after training. Interestingly, at 17-18 days after training, the increase was only observed in the context test (p<0.01, Fig. 8F), while no significant change was seen in the tone test (p>0.05, Fig. 8E).

**Figure 8.**
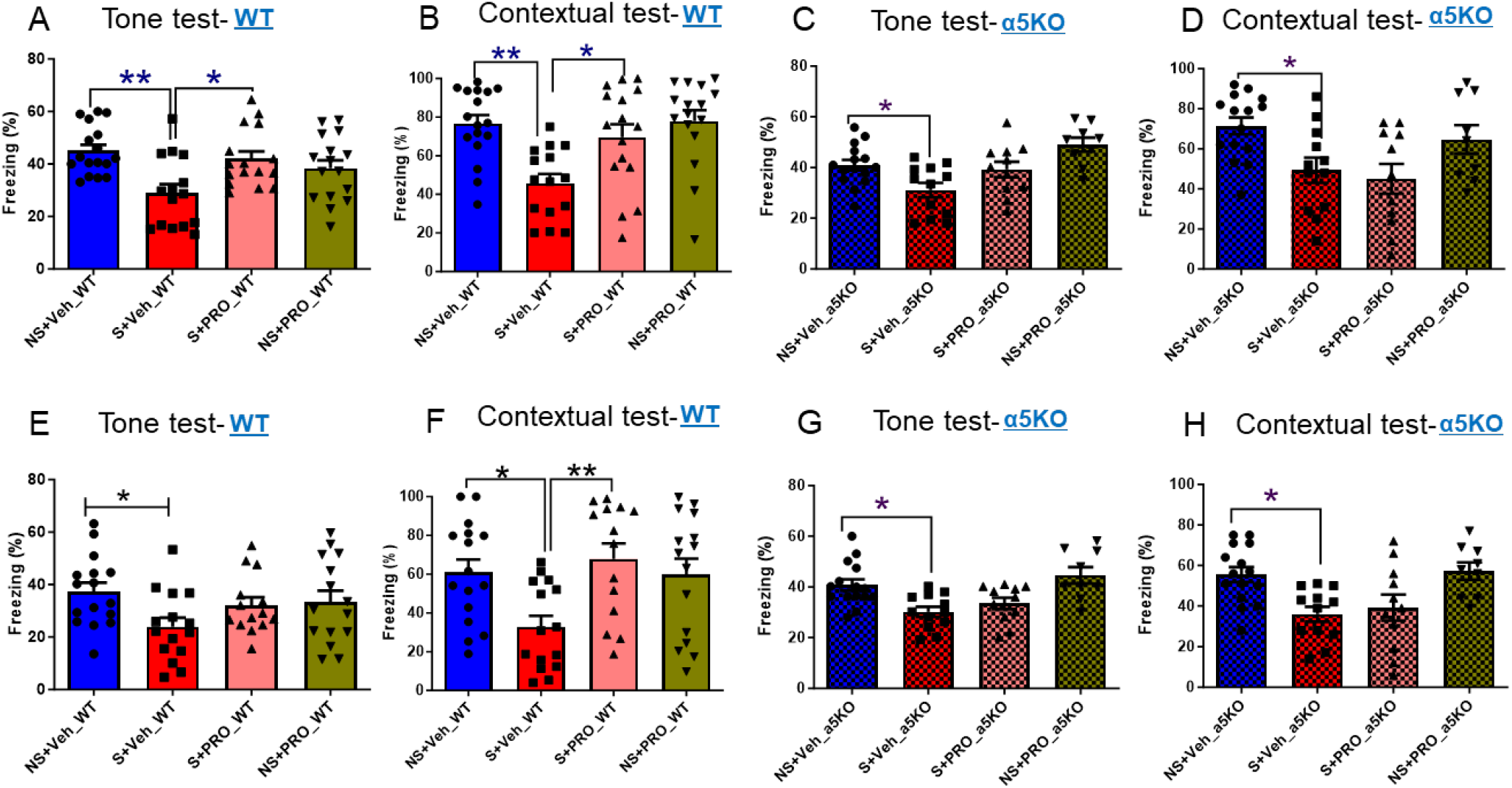
Effects of surgery and chronic intermittent propofol in trace fear conditioning and contextual conditioning in aged wild type (A, B, E, F) and α5 knockout mice (C, D, G, H). Fear conditioning was performed with recall approximately 1 week (A-D) and 4 weeks (E-H) after surgery. (**A**) Tone test for wild type mice after 1 week, F (3, 60) = 5.730, P=0.001, (comparisons: NS+Veh vs S+Veh, q=5.571; S+Veh vs S+PRO, q=4.461). (**B**) Contextual test for wild type mice after 1 week, F (3, 60) = 7.177, P=0.0003, (comparisons: NS+Veh vs S+Veh, q=5.629; S+Veh vs S+PRO, q= 4.292). (**C**) Tone test for α5 knockout mice after 1 week, F (3, 43) = 6.822, P=0.0007, (comparisons: NS+Veh vs S+Veh, q= 3.962). (**D**) Contextual test for α5 knockout mice after 1 week, F (3, 43) = 4.513, P=0.0077 (comparisons: NS+Veh vs S+Veh, q= 3.876). (**E**) Tone test for wild type mice after 4 weeks, F (3, 56) = 2.603, P=0.0609, (comparisons: NS+Veh vs S+Veh, q=3.860) (**F**) Contextual test for wild type mice after 4 weeks, F (3, 56) = 4.647, P=0.0057, (comparisons: NS+Veh vs S+Veh, q= 4.008; S+Veh vs S+PRO, q= 4.842). (**G**) Tone test for α5 knockout mice after 4 weeks, F (3, 43) = 7.125, P=0.0005, (comparisons: NS+Veh vs S+Veh, q= 4.744). (**H**) Contextual test for α5 knockout mice after 4 weeks, F (3, 43) = 5.624, P=0.0024, (comparisons: NS+Veh vs S+Veh, q= 4.552). Results are expressed as mean ± SEM and statistically evaluated by one-way analysis of variance followed by Tukey’s Multiple Comparison Test. Statistical significance: *p<0.05, **p<0.01. Abbreviations: NS+Veh: no surgery control + vehicle. S+Veh: surgery + vehicle. S+PRO: surgery + propofol. NS+PRO: no surgery control + propofol. WT: wild type mice. α5KO: α5 knockout mice.

In aged α5 KO mice, we found that surgery resulted in a reduction in freezing time in both the tone test (p<0.05, Fig 8C,G) and the context test (p<0.05, Fig 8D,H) at both 1-2 days (Fig. 8C,D) and 17-18 days (Fig. 8G,H) after training. However, treatment with CIP did not reverse the surgery-induced reduction of freezing in either the tone test (p>0.05, Fig 8C,G) or the context test (p<0.05, Fig 8D,H) at 1-2 days (Fig. 8C,D) or 17-18 days (Fig. 8G,H) after training. Notably, CIP had no effect on fear conditioning in the absence of surgery.

These findings clearly demonstrate that CIP treatment can effectively reverse the cognitive impairments induced by surgery, and that this reversal is mediated by α5 GABA_A_ receptors.

#### 2.2.6. Expression of microglial activation and proapoptotic markers

Perioperative neurocognitive disorder (NCD) (3) is associated with the inflammatory activation of hippocampal microglia(26) and with apoptosis (27). To investigate the potential involvement of microglial activation and apoptosis in surgery-induced cognitive dysfunction, we examined the expression levels of the microglial activation marker Iba-1, as well as the proapoptotic markers cleaved caspase-9, cleaved caspase-3, and Bax in the hippocampus using Western blot analysis (Fig. 9). In our study with aged wild type mice and aged α5 KO mice, we observed an increase in the expression of Iba-1 (p<0.05, Fig. 9B,G), cleaved caspase-9 (p<0.05, Fig. 9C,H), cleaved caspase-3 (p<0.05, Fig. 9D,I), and Bax (p<0.05, Fig. 9E,J) proteins following surgery compared to the non-surgery controls. Notably, this surgery-induced increase in expression was attenuated by CIP treatment in aged wild type mice, as evidenced by the significant decrease in the expression of Iba-1 (p<0.05, Fig. 9B), cleaved caspase-9 (p<0.05, Fig. 9C), cleaved caspase-3 (p<0.05, Fig. 9D), and Bax (p<0.05, Fig. 9E) compared to the surgery + vehicle group. However, CIP treatment did not show significant effects in aged α5 KO mice (p>0.05, Fig. 9G-J). Importantly, there were no significant differences between the control (no surgery control) and propofol per se (no surgery + propofol) groups in both WT mice (p>0.05, Fig. 9B-E) and α5 KO mice (p>0.05, Fig. 9G-J). These findings clearly highlight the efficacy of CIP treatment in reversing the microglial activation (Iba-1) and apoptosis (cleaved caspase-9, cleaved caspase-3, and Bax) induced by surgery, and that these effects are mediated by α5-GABA_A_ receptors. Again, in the absence of surgery CIP had no effect in either WT mice (p>0.05, Fig. 9B-E) or α5 KO mice (p>0.05, Fig. 9G-J).

**Figure 9.**
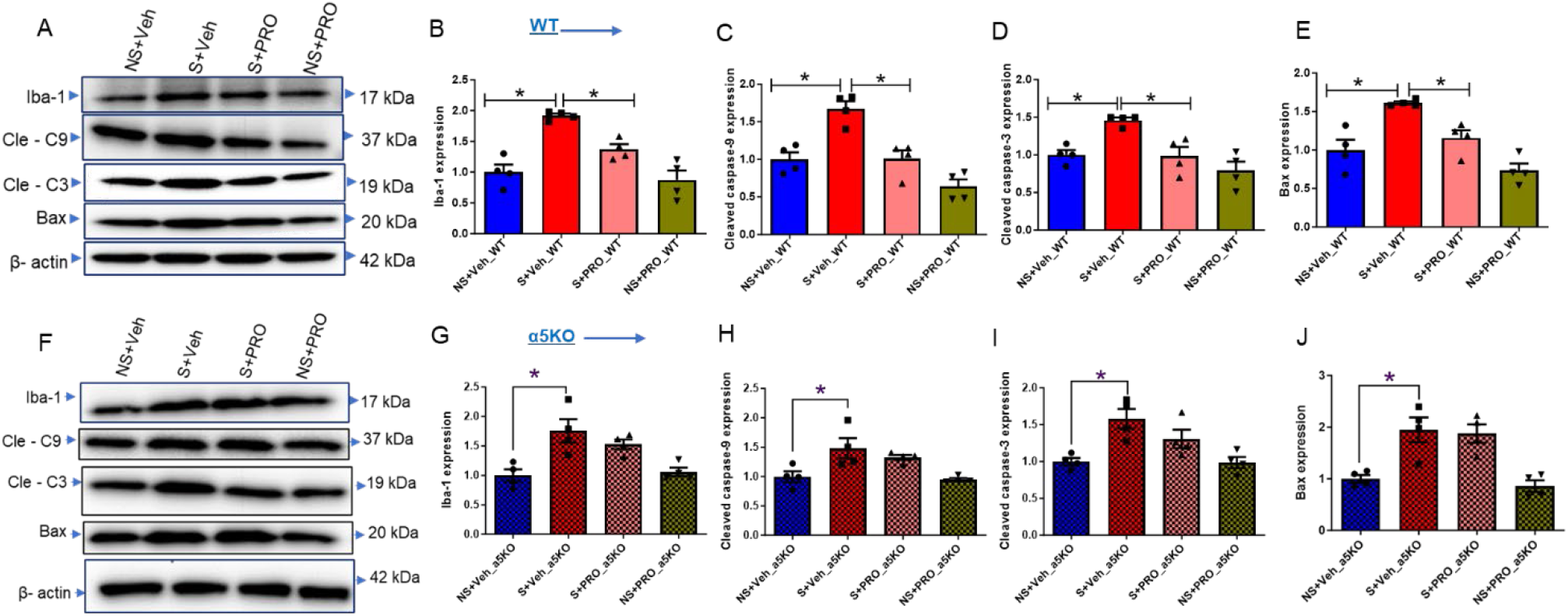
Effects of surgery and chronic intermittent propofol on microglial activation and apoptotic markers in aged wild type (A-E) and α5 knockout mice (F-J). **A**) Representative Western blot images of Iba-1, cleaved caspase-9, cleaved caspase-3, Bax, and β-actin in different groups of wild type mice. Bar graph images showing the mean ± S.E.M. of relative protein expression of **B**) Iba-1, **C**) cleaved caspase-9, **D**) cleaved caspase-3, and **E**) Bax in different groups of wild type mice. **F**) Representative Western blot images of Iba-1, cleaved caspase-9, cleaved caspase-3, Bax, and β-actin in different groups of α5 knockout mice. Bar graph images showing the mean ± S.E.M. of relative protein expression of **G**) Iba-1, **H**) cleaved caspase-9, **I**) cleaved caspase-3, and **J**) Bax in different groups of α5 knockout mice. Banding patterns shown are from one out of four experiments. The expression levels were normalized with β-actin. Statistical significance: *p<0.05. Abbreviations: NS+Veh: no surgery control + vehicle. S+Veh: surgery + vehicle. S+PRO: surgery + propofol. NS+PRO: no surgery control + propofol. WT: wild type mice. α5KO: α5 knockout mice. Cle-C9: Cleaved caspase-9. Cle-C3: Cleaved caspase-3. Statistical analyses were performed using GraphPad Prism, employing 1-way ANOVA followed by Tukey’s test for multiple comparisons. The results are as follows: B) F (3, 12) = 18.56, P<0.0001; (comparisons: NS+Veh vs S+Veh, q=8.406; S+Veh vs S+PRO, q=5.021); C) F (3, 12) = 18.67,P<0.0001; (comparisons: NS+Veh vs S+Veh, q= 6.742; S+Veh vs S+PRO, q= 6.644); D) F (3, 12) = 9.721, P=0.0016; (comparisons: NS+Veh vs S+Veh, q= 5.042; S+Veh vs S+PRO, q= 5.207); E) F (3, 12) = 14.56, P=0.0003; (comparisons: NS+Veh vs S+Veh, q= 6.328; S+Veh vs S+PRO, q= 4.720); G) F (3, 12) = 8.541, P=0.0026 (comparisons: NS+Veh vs S+Veh, q= 6.029; S+Veh vs S+PRO, q= 1.812); H) F (3, 12) = 6.496, P=0.0074 (comparisons: NS+Veh vs S+Veh, q= 4.778); I) F (3, 12) = 7.798, P=0.0038 (comparisons: NS+Veh vs S+Veh, q= 5.729); J) F (3, 12) = 12.20, P=0.0006 (comparisons: NS+Veh vs S+Veh, q= 5.805).

#### 2.2.7. Nitrite levels, a marker of oxidative stress

Finally, we assessed the effects of perioperative CIP treatment and/or surgery on nitrite levels, a marker of oxidative stress, in both aged wild type and α5 KO mice. The nitrite levels were significantly increased in the hippocampus of the surgery + vehicle group (S+Veh) compared to the no surgery + vehicle group (NS+Veh) (p<0.05, Figure 10B, C). However, CIP treatment decreased the nitrite levels in the surgery + propofol group (S+PRO) as compared to the surgery + vehicle group (S+Veh) in aged WT mice (p<0.05, Figure 10B) but not in aged α5 KO mice (p>0.05, Figure 10C). CIP had no effect in the absence of surgery (p>0.05, Figure 10B,C). Our findings indicate that surgery increases oxidative stress, that CIP reverses this oxidative stress and that this action of CIP is mediated by α5-GABA_A_ receptors.

**Figure 10.**
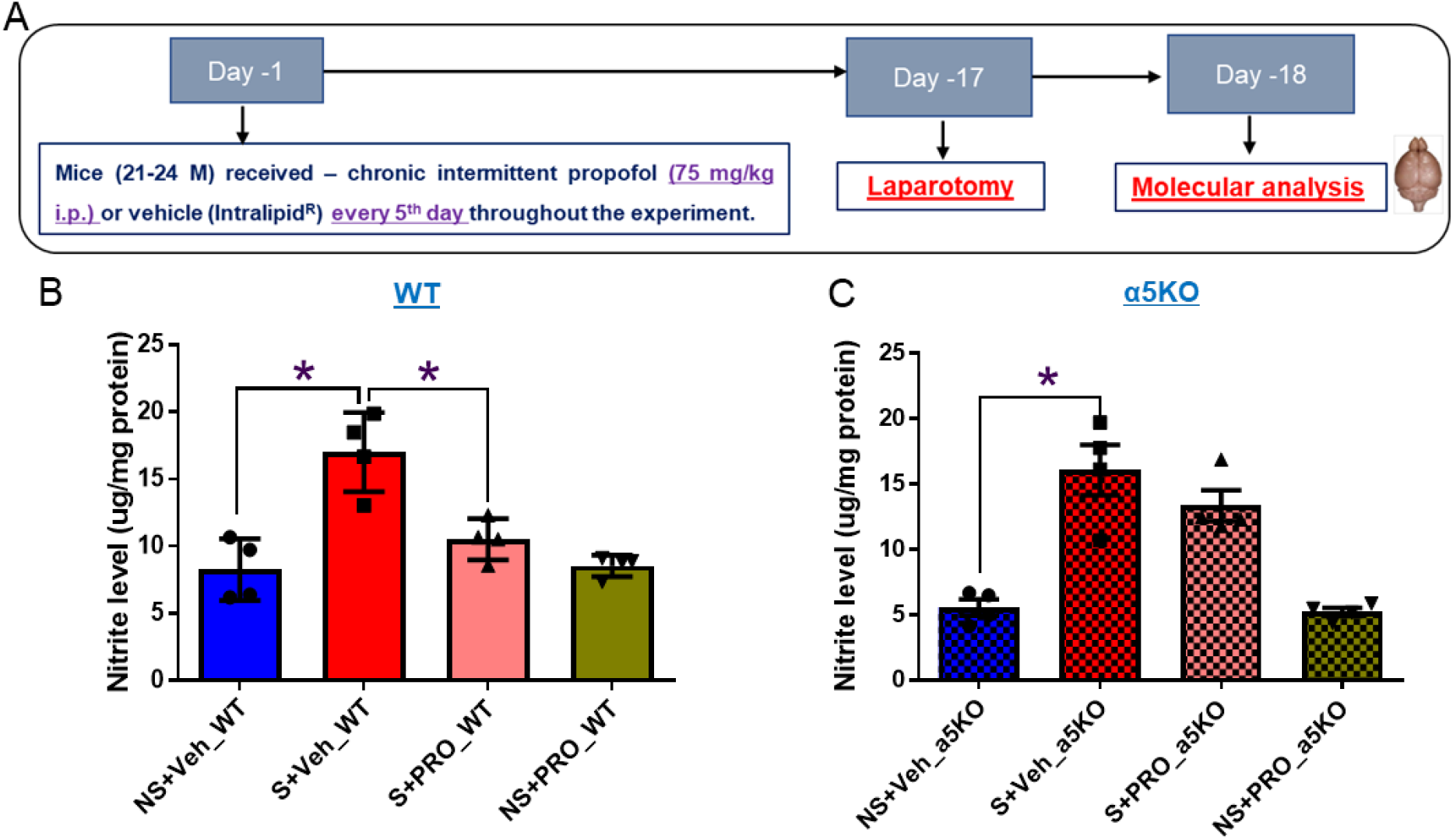
Effects of surgery and chronic intermittent propofol on oxidative stress marker (nitrite) in the hippocampus of aged wild type (B) and α5 knockout mice (C). **A**) Schematic overview of the experimental protocol for studying the effects of perioperative chronic intermittent propofol (CIP). Results are expressed as mean ± SEM and statistically evaluated by one-way analysis of variance followed by Tukey’s Multiple Comparison Test.*p<0.05. **B**) Oxidative stress marker (nitrite μg/mg protein) in wild type mice, F (3, 12) = 15.74, P=0.0002; (comparisons: NS+Veh vs S+Veh, q= 8.517; S+Veh vs S+PRO, q= 6.316). **C**) Oxidative stress marker (nitrite ug/mg protein) in α5 knockout mice, F (3, 12) = 21.47, P<0.0001; (comparisons: NS+Veh vs S+Veh, q= 8.853). Abbreviations: NS+Veh: no surgery control + vehicle. S+Veh: surgery + vehicle. S+PRO: surgery + propofol. NS+PRO: no surgery control + propofol. WT: wild type mice. α5KO: α5 knockout mice.

## 3. Discussion

Perioperative neurocognitive disorder (NCD) is a growing medical problem leading to significant morbidity in an aging population, and novel avenues for its prevention or therapy are needed. The goal of this study was to investigate whether chronic intermittent treatment with propofol is able to attenuate cognitive impairments that occur postoperatively in aged mice and, if this is the case, to determine which pathophysiological mechanisms and molecular target(s) would be responsible for this action. We found that CIP did indeed prevent the surgery-induced decline in cognitive function, that its administration resulted in the redistribution of α5-GABA_A_Rs to the cell surface of hippocampal neurons, and that the salutary effect of CIP on cognitive function was absent mice lacking α5-GABA_A_ receptors. Moreover, we found that CIP attenuated the nitrosative stress and increased expression of markers of microglial activation (Iba-1) and apoptosis (cleaved caspase-9, cleaved caspase-3, and Bax) induced by surgery, and these effects were again absent in mice lacking α5-GABA_A_ receptors. Taken together, these findings suggest a model in which CIP prevents postoperative cognitive deficits by inducing a redistribution of α5-GABA_A_ receptors to the cell surface, leading to a reduction in inflammation-induced neuronal dysfunction and cell death. However, the biochemical events that result in α5-GABA_A_R redistribution, and the signaling pathways that attenuate inflammation-related changes, remain undefined.

Propofol is a frequently used intravenous anesthetic with immobilizing, hypnotic and amnestic actions that have been shown to be mediated by GABA_A_ receptors(15, 28). It has also been reported that CIP (here 50 mg/kg propofol i.p. once per week) improves spatial memory in aged mice (20- 22 months) and in a mouse model of Alzheimer’s disease(12). In our study, abdominal surgery led to nitrosative stress, and increased expression of markers of microglial activation (Iba-1) and apoptosis (cleaved caspase-9, cleaved caspase-3, and Bax) in the hippocampus, resulting in impaired learning and memory. However, locomotor activity remained unaffected. Since CIP treatment resulted in a sustained redistribution of α5-GABA_A_ receptors to the cell surface membranes and suppression of nitrosative stress, microglial activation, and apoptosis in the hippocampus of aged mice (21-24 months of age),these changes likely underly a large part the protective role of CIP treatment against surgery-induced memory impairments. As these changes are absent in mice lacking α5-GABA_A_ receptors, our experiments show that they are mediated by α5-GABA_A_ receptors. We have thus identified a major molecular target for the prevention of neuroinflammatory, proapoptotic and cognitively impairing actions of laparotomy surgery.

Zurek et al. (2014) reported that etomidate and isoflurane lead to a redistribution of α5-GABA_A_ receptors to the cell surface membranes, which is associated with increased tonic inhibition in CA1 pyramidal neurons and impairments in the novel object recognition task, which can last for seven days (19). In separate studies, these authors reported that pretreatment with L-655,708, an α5 negative allosteric modulator, reversed the etomidate-induced or isoflurane-induced memory impairment, implying that an α5 negative allosteric modulator might be suitable to prevent postoperative cognitive deficits(20, 21). In the present study, we were interested in whether propofol also results in a sustained redistribution of α5-GABA_A_ receptors to the surface membranes and whether such a sustained redistribution might prevent or reverse perioperative neurocognitive dysfunction. We found that a single dose of propofol (100 mg/kg i.p.) increased cell-surface expression of α5-GABA_A_ receptors in the hippocampus after 24 h, 72 h, and five days, but not after seven days. Furthermore, with CIP over 21 days we observed a significant up-regulation of cell-surface expression of α5 subunits, with no evidence of tolerance development to this effect. As these results were obtained in young adult mice (3-4 months old), we confirmed that in aged mice (21-24 months old) a single propofol injection (75 mg/kg i.p.) also resulted in increased cell-surface expression of α5-GABA_A_ receptors in the hippocampus after five days. Interestingly, the total expression of α5 subunits remained unchanged at all time points in all experiments, indicating that propofol likely affects the distribution of α5-GABA_A_ receptors within neurons through an unknown mechanism, possibly involving inhibition of endocytosis. These findings suggest that propofol can cause a sustained redistribution of α5-GABA_A_ receptors in the hippocampus of both young and aged mice, which could have implications for cognitive function.

In the current study, we aimed to investigate the potential preventive or therapeutic effects of perioperative CIP treatment in mitigating surgery-induced cognitive impairments, as well as its impact on nitrosative stress markers, microglial activation, and caspase activation. To achieve this, we administered CIP treatment at a dosage of 75 mg/kg propofol intraperitoneally (i.p.) every 5th day throughout the experiment. As perioperative NCD is particularly prevalent in aged patients, we chose to do our experiments with 21-24 months old mice. Also, there is a report indicating that postoperative cognitive deficits would occur in 18 months old WT mice but not in 9 months old WT mice(7), suggesting that animals >18 months of age may be preferable for the study of such deficits. Our study revealed that abdominal surgery caused significant cognitive impairment in both aged WT and aged α5 KO mice, as evident in various behavioral tests (Y-maze, NOR, trace and contextual fear conditioning, and MWM). However, perioperative CIP treatment showed significant protective effects only in the aged WT mice, effectively mitigating surgery-induced memory impairments in all the behavioral tests. In contrast, the aged α5 KO mice did not show a protective response to CIP treatment. Despite receiving CIP, these knockout mice exhibited cognitive impairments similar to the vehicle-treated surgery group, suggesting that CIP’s effectiveness in mitigating cognitive decline after surgery is critically dependent on α5-GABA_A_ receptors.

Our current study’s findings on CIP’s effects in aged WT mice are consistent with previous observations by Shao et al. (2014), who demonstrated that the effects of CIP administration (50 mg/kg/week i.p.) improved cognitive function in the water maze and attenuated Aβ-induced mitochondrial dysfunction and brain caspase-3 and caspase-9 activation in both aged wild type and AD-Tg mice (12). The propofol-induced reduction of Aβ-induced caspase-3 activation and the reduction of Aβ-induced opening of the mitochondrial permeability transition pore (mPTP) were attenuated by flumazenil, which is an antagonist at the benzodiazepine binding site of GABA_A_ receptors(29). This effect was difficult to interpret until it was reported that there are shared structural mechanisms of binding of general anesthetics and benzodiazepines to GABA_A_ receptors, providing a potential mechanism for anesthetic reversal by flumazenil (30). Limon et al. (2012) reported that patients with Alzheimer’s disease (AD) had decreased numbers of GABA_A_ receptors (including a decreased level of α5 subunit mRNA), an age-dependent reduction of GABA currents, a faster rate of desensitization and less sensitivity to GABA(31), suggesting deficits in GABA_A_-mediated neurotransmission in AD. Moreover, studies of human postmortem brain revealed that the expression of the α5 subunit is significantly decreased with age(32). In the current study, CIP treatment results in increased surface expression of α5-GABA_A_ receptors, which may underly the improvement of cognitive function in aged animals. Recently, it has been shown that propofol, administered 30 min after fear conditioning, led to reduced contextual freezing 2, 4, and 6 weeks after training, and restored shock-induced spatial memory deficits in the MWM, which is somewhat reminiscent of our findings; while the authors showed that propofol blocks shock-induced reductions of LTP and LTD, the mechanisms behind these observations are not entirely clear(33).

Numerous studies have shown that microglia are required for postoperative hippocampal inflammation and cognitive decline in mice, and that microglial activation is a key factor in the pathophysiology of perioperative NCD(26, 34). Systemic inflammation caused by surgery could induce neuroinflammation, mainly through destroying the permeability of the blood-brain barrier (35), hence, promoting the activation of microglia. Activated microglia subsequently release more inflammatory cytokines, NO and trigger apoptosis and mitochondrial dysfunction (36, 37). Excessive NO production is associated with cellular modifications, including microglial activation, neuronal cell apoptosis, and oxidative stress, leading to impaired cognitive function. The bioavailability of NO is considered a predictive risk factor for Alzheimer’s disease (AD) and early postoperative cognitive dysfunction (27, 38). In our study, we observed an increase in the activated microglial marker Iba-1 in both aged wild-type and α5 knockout mice after abdominal surgery. This increase was accompanied by elevated NO generation, and upregulation of pro-apoptotic markers, including cleaved caspase-9, cleaved caspase-3, and Bax in the hippocampus. These findings suggest a significant inflammatory and apoptotic response in the hippocampus following laparotomy surgery. However, the administration of CIP treatment showed differential effects in the two groups of mice. In aged wild-type mice, CIP treatment effectively attenuated NO generation, and the expression of activated microglia and pro-apoptotic markers, providing protection against surgery-induced memory impairment. In contrast, the protective effects of CIP were not observed in the α5 KO mice, indicating that α5-GABA_A_ receptors are critical for the efficacy of CIP in mitigating cognitive decline following surgery in WT mice. These findings align with a report where CIP administration attenuated Aβ-induced caspase activation and Aβ-induced mPTP opening in a flumazenil-sensitive manner and thus likely via GABA_A_ receptors(12). The precise mechanism for this is unclear but could involve upstream radical scavenging (39) or a direct cellular effect on cytokine responsiveness or secretion, or both. For example, propofol completely ablated lipopolysaccharide-stimulated cytokine secretion in cultured microglial cells, in contrast to volatile anesthetics(40, 41). Perioperative depletion of microglia attenuates surgery-induced cognitive decline in mice (26). Furthermore, in humans undergoing anesthesia with propofol reduced gene expression of pro-inflammatory cytokines was found in macrophages (42). Taken together, these findings suggest a potential association between GABAergic signaling and caspase activation, neuroinflammation, and cognitive function. Future studies may use a different anesthetic agent, e.g., etomidate to test this hypothesis further. These findings may promote more research leading to new concepts of perioperative NCD neuropathogenesis and new intervention (s) for perioperative NCD.

As mentioned previously, it has been suggested previously that an α5 negative allosteric modulator would be suitable for prevention or treatment of postanesthetic and thus potentially postoperative cognitive deficits(20, 21), which appears to be in contrast to our finding that α5-GABA_A_ receptors mediate a reversal of postoperative cognitive impairments. It is noteworthy that initial results with α5 knockout mice indicated that these mice display a shorter latency to find the hidden platform in the MWM(17), indicating that α5-GABA_A_ receptors would constrain spatial memory(43), and thus an α5 negative allosteric modulator would be expected to improve memory. The experiments cited above were performed in young adults, but not in aged mice. There is now evidence that in aged rodents α5 positive allosteric modulation improves cognition (22, 44), likely due to an age-related deficiency of the α5-GABA_A_ receptor system. Interestingly, we observed that CIP increased the performance in the Y maze and the NOR test in the groups not receiving surgery, consistent with these cited reports using aged mice. However, in all other behavioral tests, CIP had no effect in the absence of surgery. This suggests that the effects of CIP largely do not represent a “correction” of an age-related deficiency in the α5-GABA_A_ receptor system, but a prevention or reversal of the surgery-induced neuroinflammation. It is our interpretation that the previous observation by others that an α5 negative allosteric modulator improves general anesthetic-induced cognitive impairments is dependent on the use of young adult (and not aged) mice in their studies(20, 21), and that their recommendation for α5 negative allosteric modulators to treat general anesthetic-induced cognitive deficits may only apply to experiments with young adult animals and experimental setups not including surgical interventions.

Our studies have several limitations. First, we did not determine the dose-or time-dependent effects of propofol on surgery-induced memory impairment in aged mice, in part due to logistic constraints. Since in the literature propofol showed dual effects, i.e., neuroprotection(12, 41) and neurotoxicity(14, 45) in the brain, it is possible that propofol treatment with different doses or administered at different times may have different effects. Nevertheless, our current studies have demonstrated that propofol induces a sustained redistribution of α5-GABA_A_ receptors to the cell surface membranes in the hippocampus and validated the concept that CIP can indeed prevent or reverse perioperative NCD.

Second, we did not directly assess the role and the cellular location of α5-containing α5-GABA_A_ receptors mediating the memory-enhancing effect of CIP on surgery-induced memory impairment in aged mice. Future studies could evaluate whether the cognition-enhancing effect is mediated by α5-GABA_A_ receptors in hippocampal principal or other neurons. For example, recently, it was shown that the amnestic effect of etomidate is mediated by a5-GABA_A_ receptors on hippocampal interneurons and not on principal neurons (46). It is also possible that the effects of propofol observed in our studies is mediated by α5-GABA_A_ receptors on microglia cells. However, the main objective of the current studies was to determine whether CIP results in a reduction in perioperative NCD in aged mice and to determine the molecular target mediating this effect. Future studies will systematically investigate the underlying cellular mechanism by which CIP improves memory function.

In conclusion, our study demonstrates that chronic intermittent propofol treatment improves cognitive function postoperatively in aged wild-type mice, and that most likely a sustained redistribution of α5-GABA_A_ receptors to the cell surface membranes underlies these effects. This treatment also effectively attenuates the surgery-induced increase in nitrosative stress, and pro-apoptotic proteins (caspase-9, cleaved caspase-3, and Bax), as well as microglial cell activation in the hippocampus. However, these beneficial effects were not observed in the α5 KO mice. These findings suggest that CIP treatment may have a preventive or therapeutic role in mitigating surgery-induced cognitive dysfunction and reducing the risk of long-term neurocognitive outcomes. The increased availability of α5-GABA_A_ receptors on cell surface membranes after CIP treatment appears to be an important mechanism underlying its cognitive protective effects in the context of surgery-induced cognitive impairment in aged mice. It will be interesting to evaluate whether other modulators of the GABA_A_ receptor system (e.g., other general anesthetics, benzodiazepines, neurosteroids) have effects similar to those of CIP. Furthermore, the perioperative use of subtype-selective α5 positive allosteric modulators(47) could be a promising avenue for the development of a relatively simple and widely applicable clinical protocol to mitigate or prevent perioperative NCD in elderly patients.

## 4. Methods

### Animals

All experiments were conducted under oversight and with the approval of the Institutional Animal Care and Use Committee (IACUC) of the University of Illinois Urbana-Champaign. C57BL/6J mice (Jackson Laboratory stock# 000664) were purchased at 1-2 months of age and maintained under controlled laboratory conditions (temperature: 22 ± 2°C; humidity: 55 ± 5%; light on from 0700-1900). Food and water were available ad libitum. All procedures were consistent with the Guide for the Care and Use of Laboratory Animals of National Research Council, and all of efforts were made to minimize the number of animals in the studies. ARRIVE guidelines were followed. Animals were randomized into experimental groups and research was performed blinded. All experiments were approved by the University of Illinois Urbana-Champaign Institutional Animal Care and Use Committee (IACUC,).

### Generation and breeding of global α5 knockout mice

The generation of global α5 knockout (α5 KO) mice (Gabra5^tm2.2Uru^) has been reported previously (48). The α5 KO allele was backcrossed to C57BL/6J mice obtained from The Jackson Laboratory (strain #000664). This backcrossing process was conducted for a minimum of nine generations to ensure the desired genetic background. To verify the genotype of the resulting offspring, genotyping was carried out using DNA templates obtained from tail tips. PCR primers, namely P1 (TTTAGTGTGGGTGGTGATAGGT), P2 (CTTCCACAACGGCAAGAAGTCC), and P3 (CCACAGATACCCAGATGAATGTG), were employed.

### Cell-surface biotinylation

Hippocampal tissue from 3-4 months old C57BL/6J mice was used for these experiments. Coronal hippocampal slices (1 mm) were prepared 24 h, 72 h, 5 days, or 7 days after treatment with propofol (PropoFlo^TM^28, Zoetis, USA) (100 mg/kg, i.p) or chronic intermittent propofol (CIP, 75 mg/kg, i.p., every 5^th^ day for 21 days) or vehicle (Intralipid^R^) and placed in oxygenated aCSF (95% O_2_, 5% CO_2_) for 30 min to recover. Slices were then placed on ice and incubated twice with 0.75 mg/ml NHS-SS-biotin in PBS (Thermo Scientific, Rockford, Illinois) for 30 min while bubbling with 95% O_2_ and 5% CO_2_. Excess biotin was quenched with a quenching solution (Pierce cell surface isolation kit, Thermo Scientific) and removed by washing slices three times with ice-cold aCSF. Slices were then lysed with a mild detergent, and the labeled proteins were then isolated with Thermo Scientific™ NeutrAvidin™ Agarose. The bound proteins were released by incubating with an SDS-PAGE sample buffer containing 50mM DTT. Proteins were separated by sodium dodecyl sulfate-polyacrylamide gel electrophoresis (SDS-PAGE), transferred onto polyvinylidene fluoride (PVDF) membranes, and incubated at 4◦C overnight with primary antibodies for the α5-GABA_A_ receptor (1:2, NeuroMab clone N415/24), Na+/K+ ATPase (1:200, Santa Cruz Biotechnology, Catalog # SC-514661) and β-actin (1:2000, Cell Signaling Technology, Catalog # 4970). The membranes were washed with TBST three times and then incubated at room temperature for 60 min with secondary antibody goat anti-rabbit IgG (1:3000, Cell Signaling Technology, Catalog # 7074) or goat anti-mouse IgG (1:3000, Cell Signaling Technology, Catalog # 7076). Then, the membranes were washed with TBST four times. The SuperSignal West Pico Chemiluminescent Substrate kit from Thermo Fischer Scientific (Rockford, IL) was used to detect antigens utilizing the manufacturer’s instructions.

To further investigate the age-dependent response of propofol on the cell surface expression of α5- GABA_A_ receptors in the hippocampus, cell-surface biotinylation experiments were conducted in 21-24-month-old C57BL/6 J mice. These mice received a single dose of propofol (PropoFlo^TM^28, Zoetis, USA) (75 mg/kg, i.p) or vehicle (Intralipid^R^), and biotinylation was performed five days after the administration. Subsequently, Western blot analysis was conducted to examine the expression of α5-GABA_A_ receptors.

### Experimental protocol and administration of Propofol

To study the effect of chronic intermittent propofol (CIP) on surgery-induced memory impairment in 21-24 months old wild type C57BL/6J mice, a dose of 75 mg/kg, i.p. every 5^th^ day, was selected based on an initial dose-finding study. Vehicle groups received the same volume of Intralipid^R^ (i.p.) every 5^th^ day throughout the experiment. After propofol treatment the mice were placed in a heated (37◦C) and humidified chamber flushed continuously with oxygen (100%), and continuously observed for breathing, heart rate, and skin color for approximately 30-60 minutes.

Aged wild type (WT) or α5 knockout (α5 KO) mice, aged between 21 to 24 months, were randomly assigned to four groups. Each group received a different treatment throughout the experiment:

Group 1 : No surgery + Vehicle- (Intralipid^R^), every 5^th^ day throughout the experiment.
Group 2 : No Surgery + Propofol- (75 mg/kg, i.p)- every 5^th^ day throughout the experiment
Group 3 : Surgery + Vehicle- (Intralipid^R^), every 5^th^ day throughout the experiment
Group 4 : Surgery + Propofol- (75 mg/kg, i.p), every 5^th^ day throughout the experiment

### Laparotomy

We performed a laparotomy in aged WT and α5 KO (21-24 months old) mice under 10 min isoflurane (2%) anesthesia. A 2.5 cm incision was made in the middle of the abdomen to open the abdominal cavity, and the abdominal cavity and skin were sutured and closed. All animals received 0.1 mg/kg buprenorphine s.c. immediately before the surgery for pain control. Mice were kept warm on a heating pad as they recovered. Mouse chow moistened with water was placed in the cage once they recovered to encourage eating and provide hydration following surgery. The cages were kept on heating pads as necessary the day following surgery.

### Open field test

Open field tests were used to evaluate the locomotor and exploratory activities of mice in a novel environment. An open field box (40 cm × 40 cm × 30 cm) was made from plexiglass. Animals were placed randomly in the center of the open field box, and their activity was observed for 10 minutes using the EthoVision XT video tracking system, which was mounted above the chamber. The open field chamber was wiped with 70 % ethanol before use and subsequent tests to remove any scent clues left by the previous subject mouse. The total distance traveled (cm) and time spent in the inner zone of the open field chamber was observed.

### Spontaneous alternation Y-maze test

Spontaneous alternation is a measure of spatial working memory. Each animal was placed in the center of a symmetrical Y-maze and was allowed to explore freely during an 8-min session. The sequence and total number of arms entered were recorded with a video camera that was mounted above the apparatus. Arm entry was defined as entry of the whole body into an arm. Alternation was defined as successive entries (ABC, ACB, BCA, BAC, CBA, CAB) into the three arms in overlapping triple sets. The percentage of correct alternations was calculated as the number of triads containing entries into all three arms divided by the maximum possible number of alternations (the total number of arms entered − 1) × 100. To diminish odor cues, the Y maze was cleaned with 70% ethanol between trials and allowed to dry.

### Novel object recognition test

Object recognition was assessed in a 40 cm × 40 cm × 30 cm opaque chamber in a dimly lit room. Mice are habituated to the chamber for two days, 10 min per day. During the training phase, on day three, the mouse was allowed to explore two copies of the same object for 10 min. The mouse was then returned to its home cage for a retention period of 1 h. The mouse was reintroduced to the training context and presented with one familiar sample object and one novel object for 10 min. Movement and interaction with the objects were recorded and analyzed with a videocamera which was mounted above the chamber and connected the EthoVision XT video tracking system. Exploratory behavior was measured. Exploratory behavior was defined as sniffing, licking, or touching the object while facing the object. Interaction time was recorded using the multiple body point module. Memory was assessed by determining the recognition index (i.e., the ratio of time spent exploring the novel object to the time spent exploring both objects). After each mouse had performed the test, the chamber and objects were swabbed thoroughly with alcohol to avoid any interference with tests involving subsequent mice.

### Trace -and Contextual Fear conditioning

On the training day of the experiment, the mice were placed in a conditioning box (Med-Associates, Inc., St. Albans, VT) with grid floor. They were allowed to explore the fear conditioning chamber for 180 seconds before presented with a 2-Hz pulsating tone (20 sec,70 dB, 5000 Hz) - shock (2 sec, 0.7 mA) sequence separated by an empty 20 sec trace interval. The pairing of stimuli was repeated five times at 1-min intervals. Sixty seconds after the fifth shock, the mice were removed from the chamber. The first contextual test was performed at ∼ 24 hours after the end of the pairing. Each mouse was allowed to stay in the same training chamber for the contextual test for a total of 3 min, with no tones or shocks delivered, and freezing behavior was recorded. At the end of the contextual test, the mice were returned to their home cage. After ∼ 24 hours, mice were placed in a novel environment for a cued fear memory test for a total of 9 min. The new environment consisted of a colored Plexiglas sheet that covered the steel rods of the floor, black striped plastic on the chamber’s walls, and the introduction into the testing chamber of a novel odor (1% acetic acid as olfactory cue). After 3 min without any stimulus, mice were exposed to the auditory cue for the remaining minutes (6 min), and freezing behavior was scored. Cognitive function in the tone test was also assessed by measuring the freezing time. The same cohorts of mice were also tested in the context and tone testing at four weeks post-surgery without additional pairing.

### Morris Water Maze and Reversal Learning

The spatial learning and memory function of mice were assessed using the Morris water maze. The test lasted for 14 days: learning phase sessions 1- 8 (day1-8); learning phase probe trials (Day 9); reversal learning phase sessions 1-4 (day 10-13) and reversal learning probe trials (day 14). Mice are tested in a pool (120 cm in diameter) filled with water (23-25°C) made opaque with the addition of a white nontoxic dye (Premium Grade Tempera, Blick, Galesburg, IL) containing a platform (10 cm in diameter) that is submerged by 1 cm under the water surface. Geometric shapes are affixed to the walls to serve as extra-maze cues. Mice are given 3 trials every day, being released from a different quadrant each time in random order, with the platform location constant. A trial ends either 2 sec after the mouse climbs on the platform, or 60 sec after the start of the trial, with the experimenter guiding the mouse to the platform. If a mouse could not find the platform within 60 sec it was gently guided to the platform and the escape latency was recorded as 60 sec. After the mouse reached the platform, it was allowed to stay there for 10 sec before the next test was performed. From day 10 to day 13, the reversal learning phase was established by moving the platform from the original location to the nearest quadrant to increase the effects of interference. During probe trial (day 9) and reversal learning probe trial (day 14), the platform was removed, and the mice were left in the pool for 60 seconds. The track of mice was recorded using a camera positioned above the center of the pool and connected to the EthoVision XT software video tracking system.

### Biochemical and molecular parameters

Biochemical and molecular parameters estimations were performed in the treatment group. For biochemical assays, animals were deeply anesthetized with ketamine (139 mg/kg i.p.) / xylazine (21 mg/kg i.p.) and then transcardially perfused with ice-cold phosphate-buffered saline (PBS). The brain was removed quickly, kept on an ice-cold plate, and then dissected to harvest the hippocampus.

### Western blots

Hippocampal tissue samples were homogenized using RIPA Lysis and Extraction Buffer (Thermo Fisher Scientific, Catalog number: 89900), which was supplemented with protease inhibitors and phosphatase inhibitors. The homogenate was centrifuged at 13,000 rpm for 30 min at 4°C, and supernatant was collected for measurement of protein concentrations with a BCA assay kit. Proteins were separated by sodium dodecyl sulfate polyacrylamide gel electrophoresis (SDS-PAGE), transferred onto polyvinylidene fluoride (PVDF) membranes and incubated at 4°C overnight with primary antibodies for Cleaved Caspase-3 (Asp175) (5A1E) [1:1000, Cell Signaling Technology, catalog # 9664], Cleaved Caspase-9 (Asp353) [1:1000, Cell Signaling Technology, catalog #9509], Bax [1:1000, Cell Signaling Technology catalog #14796], Iba-1 [1:500, Invitrogen, catalog # MA5-36257] and β-actin [1:2,000, Cell Signaling Technology, catalog # 4970]. The membranes were washed with TBST three times and then incubated at room temperature for 60 min with secondary antibody goat anti-rabbit IgG (1:3000, Cell Signaling Technology, catalog # 7074) or goat anti-mouse IgG (1:3,000, Cell Signaling Technology, catalog # 7076). Then the membranes were washed with TBST four times. The SuperSignal West Pico Chemiluminescent Substrate kit from Thermo Fischer Scientific (Rockford, IL) was used to detect antigens utilizing the manufacturer’s instructions.

### Nitrite estimation

Mice were deeply anesthetized with ketamine (139 mg/kg i.p.) / xylazine (21 mg/kg i.p.) and then transcardially perfused with ice-cold normal saline. After perfusion, the brains were harvested from the mice. Nitrite levels, serving as an indicator of nitric oxide (NO) levels, were estimated in the hippocampus from various experimental groups using the Griess reagent(0.1% *N*-(1-naphthyl) ethylenediamine dihydrochloride, 1% sulfanilamide, and 2.5% phosphoric acid. Sodium nitrite was employed as a standard. Equal volume of Griess reagent and supernatant from different regions of brain homogenate were incubated at 37 °C for 30 min followed by absorbance reading at 542 nm wavelengths spectrophotometrically in ELISA reader (Biotek). It was expressed as μg/mg protein.

### Statistical analysis

Statistical analyses were performed (GraphPad Prism) using 1-way ANOVA followed by Tukey’s test for multiple comparisons or Two-way ANOVA followed by Sidak’s multiple comparisons test for multiple comparisons or two-tailed t-test. The results are presented as mean ± standard error of the mean (SEM), and the p-values are indicated in each figure legend. Statistical significance was defined as p < 0.05.

## Author contributions

Conceptualization: UR; Investigation: RN; Analysis and Interpretation: RN, JL, MK, UR; Methodology: RN, JL, MK, MW, RAP, UR; Funding acquisition: UR; Supervision: UR; Writing – original draft: RN, UR; Writing – review & editing: RN, JL, MK, MW, RAP, UR.

## Data availability

All original research data will be archived in the Harvard Dataverse repository.

## Ethics approval

Permission for animal experiments was obtained from the University of Illinois Urbana-Champaign Institutional Animal Care and Use Committee (IACUC).

